# A novel proliferative candidate genes panel for idiopathic pulmonary fibrosis: insights from integrated bulk and single-cell RNA sequencing

**DOI:** 10.64898/2026.02.19.706933

**Authors:** Qing Wang, Chunli Tang, Qi Wu, Nansheng Wan, Zhixian Jin, Chao yang, Huiming Wang, Jing Feng, Yubao Wang

## Abstract

**Background:** Idiopathic pulmonary fibrosis (IPF) remains a fatal interstitial lung disease with limited diagnostic specificity and therapeutic options. This study integrates bulk and single-cell RNA sequencing (RNA-seq) to identify novel biomarkers and elucidate molecular mechanisms underlying IPF pathogenesis.

**Methods:** We prospectively enrolled 14 treatment-naive IPF patients and 6 controls. Bulk RNA-seq was performed on bronchoalveolar lavage fluid (BALF), while single-cell RNA-seq analyzed lung tissues from 4 IPF patients and 3 controls. Differentially expressed genes (DEGs) were identified (|log2FC| >1, FDR <0.05), followed by functional enrichment, protein-protein interaction (PPI) network analysis, and cell-type-specific expression profiling.

**Results:** 1. DEG Identification: Bulk RNA-seq revealed 108 DEGs (24 upregulated, 84 downregulated). KEGG enrichment analysis of DEGs revealed that upregulated genes were mainly enriched in inflammation and immune pathways (such as NF-κB signaling pathway, Fc epsilon RI signaling pathway, B cell receptor signaling pathway, phagosome, Fc gamma R-mediated phagocytosis), pyrimidine metabolism, cell cycle, and PI3K-Akt signaling pathway. 2. PPI Network: Module analysis identified a proliferative gene module 1 (NUF2, CEP55, ANLN, TTK, TK1, MYBL2, CCNA2, RRM2, CDT1) linked to cell division and cycle regulation. 3. Single-Cell Insights: scRNA-seq of 30,477 cells delineated 11 populations. Module 1 genes exhibited predominant expression in proliferating cells, Module 1 signature score of proliferating cells was significantly higher in IPF than in control group. 4. Pathogenic Links: Key genes (e.g., CEP55, TTK) were associated with PI3K/AKT signaling, epithelial-mesenchymal transition (EMT), and anti-apoptotic pathways, mirroring oncogenic mechanisms.

**Conclusion:** This multi-omics approach uncovers a proliferation-centric gene module in IPF, revealing shared molecular pathways with tumorigenesis. Our findings highlight novel diagnostic biomarkers and suggest repurposing cell cycle inhibitors as potential therapies. Future studies should validate these targets in preclinical models to advance precision medicine for IPF.

## 1 Introduction

Idiopathic pulmonary fibrosis (IPF) is a chronic, progressive, and fatal interstitial lung disease (ILD) with a median survival of 3–5 years following diagnosis ^[1].^ Characterized by aberrant lung fibrogenesis and histopathological patterns consistent with usual interstitial pneumonia (UIP), IPF manifests with debilitating symptoms—including persistent cough and exertional dyspnea—that profoundly impair health-related quality of life. Diagnosis requires a two-pronged approach: (1) multidisciplinary exclusion of known ILD etiologies through clinical-radiological-pathological correlation, and (2) confirmation of UIP on high-resolution computed tomography (HRCT) or surgical lung biopsy (SLB) / transbronchial lung cryobiopsy (TBLC) . TBLC was regarded as an acceptable alternative to SLB for making a histopathological diagnosis in patients with ILD of undetermined type in medical centers with experience performing and interpreting TBLC (conditional recommendation) [1].

However, diagnostic accuracy remains constrained by two critical limitations:

1. **Radiological ambiguity**: HRCT demonstrates suboptimal specificity in differentiating IPF from other fibrotic ILDs (e.g., chronic hypersensitivity pneumonitis [cHP], connective tissue disease-associated ILD [CTD-ILD]), which may exhibit radiologically overlapping UIP-like patterns ^[2,3]^.
2. **Procedural risks**: Although SLB is diagnostic gold standard, its invasiveness precludes use in patients with advanced disease or comorbidities due to high morbidity (e.g., pneumothorax, acute exacerbation) and mortality rates (3–6%) ^[1]^. Existing biomarkers (e.g., Krebs von den Lungen-6 [KL-6], surfactant protein-A/-D [SP-A/D]) exhibit limited specificity ^[4,5]^. These gaps underscore the urgent need for novel biomarkers with high diagnostic accuracy.

### Pathobiological insights

IPF has undergone a paradigm shift from initial attributions to chronic inflammatory processes [6] toward recognition of the significant role played by oxidative stress [7]. Contemporary models posit that repetitive microinjuries to the alveolar epithelium induce cellular dysfunction, triggering a self-perpetuating cascade of fibroblast activation, proliferation, and extracellular matrix (ECM) overproduction through dysregulated pro-fibrotic signaling ^[8,9]^. This maladaptive repair process culminates in irreversible parenchymal scarring and progressive loss of pulmonary function. ^[10,11]^ Despite the efficacy of antifibrotic therapies (pirfenidone, nintedanib) in slowing forced vital capacity (FVC) decline, IPF remains incurable ^[1]^. These limitations highlight the imperative to elucidate novel molecular mechanisms and therapeutic targets.

### Rationale for the present study

Bronchoalveolar lavage fluid (BALF), a minimally invasive biospecimen, offers a unique window into the alveolar microenvironment. When combined with bulk and single-cell RNA sequencing (RNA-seq), BALF enables high-resolution mapping of disease-specific transcriptomic signatures. While bulk RNA-seq identifies population-level expression trends, scRNA-seq resolves cellular heterogeneity, thereby overcoming the limitations of traditional tissue-based approaches.

**Study hypothesis**: Integrative analysis of bulk and scRNA-seq data from BALF and lung tissue will identify novel IPF biomarkers with diagnostic potential.

## 2. Methods (Technical Route Outlined in Figure 1)

**Figure 1.**
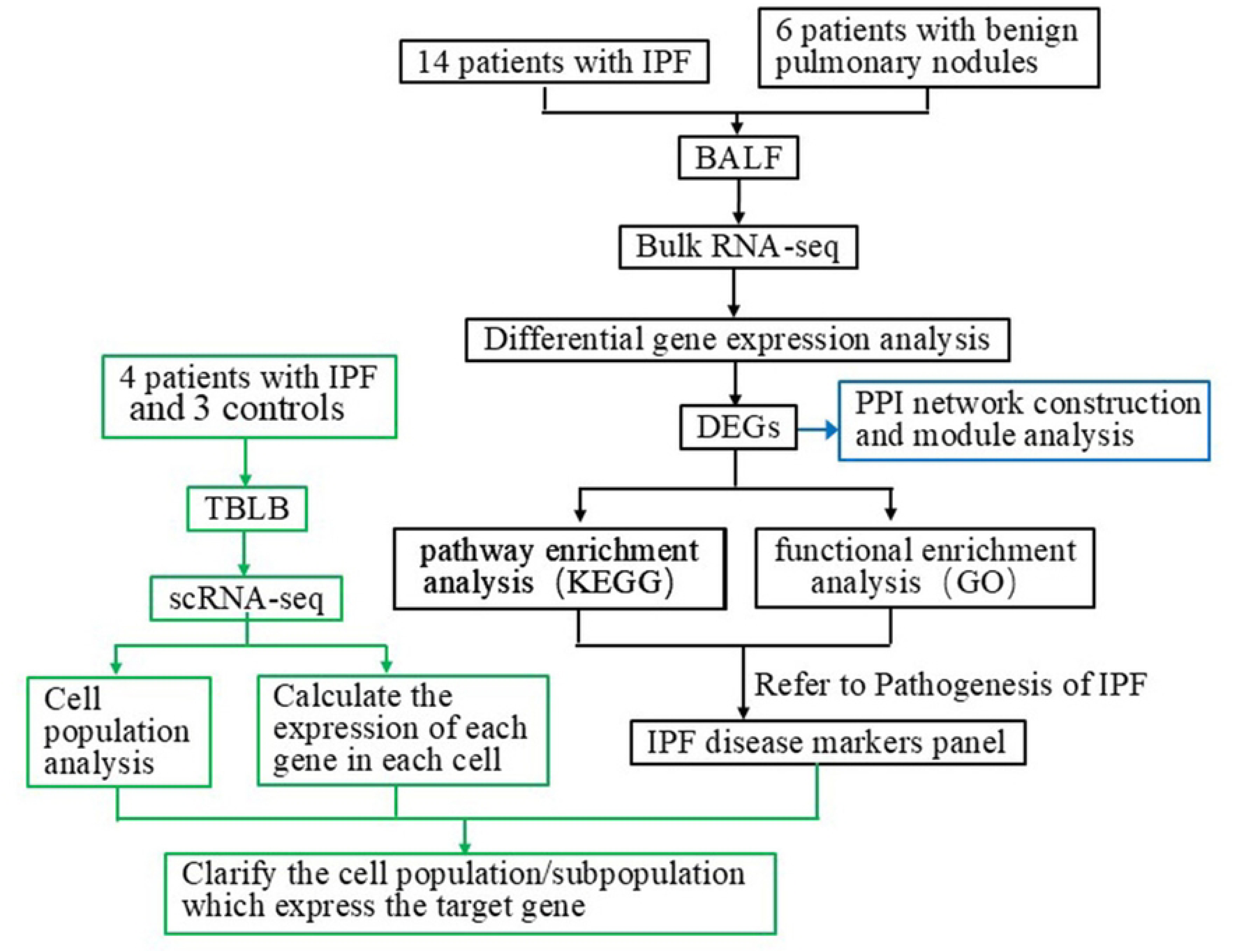
Technical Route Outlined **Abbreviations:** BALF : bronchoalveolar lavage fluid; bulk RNA-seq: bulk RNA sequencing; TBLB: transbronchial lung biopsies; DEGs: Differentially Expressed Genes.

### 2.1 Study Cohort

We prospectively enrolled 14 treatment-naive IPF patients (2023–2024) and 6 age-/sex-matched controls with benign pulmonary nodules from Kunming First People’s Hospital. Inclusion criteria:

- IPF diagnosis based on ATS/ERS/JRS/ALAT guidelines (an Update) ^[1]^.
- Exclusion of alternative ILDs via multidisciplinary discussion.

#### Data collection

- **Demographics**: Age, sex, smoking history (pack-years).
- **Radiological**: HRCT pattern: UIP, Probable UIP, Indeterminate, Alternative diagnosis.
- **Pulmonary function**: FVC, FEV₁, DLCO (predicted %), TLC.
- **Laboratory**: BALF differential counts, serum KL-6, C-reactive protein [CRP], lactate dehydrogenase [LDH], the autoantibody panel((ENA profile, ANCA, and rheumatoid-associated antibodies).
- **Ethics**: Approved by Kunming First People’s Hospital IRB (No. YLS2022-82). All participants provided written informed consent.

### 2.2 Bulk RNA-Seq and Bioinformatics

- **Sample processing**: BALF cells from IPF (n=14) and controls (n=6) underwent poly(A)+ RNA-seq (Illumina NovaSeq 6000, 150 bp paired-end).
- **Differential expression**: Differentially Expressed Genes (DEGs) were were screened using False Discovery Rate (FDR) and log2FC, with FDR < 0.05 and |log2FC| > 1. Genes with FC (i.e., fold change) greater than 2 were defined as DEGs, and genes with Log2FC>1, i.e., FC greater than 2, were considered up-regulated, and those with Log2FC<-1, i.e., FC less than 0.5, were considered down-regulated.
- **Functional enrichment**: Kyoto Encyclopedia of Genes and Genomes (KEGG) and Gene Ontology (GO) analysis (clusterProfiler, R).

### 2.3 PPI Network and Module Analysis

- **Network construction**: STRING database (confidence score >0.7).
- **Module identification**: MCODE plug-in (Cytoscape) with parameter settings: degree cutoff=2, node score cutoff=0.2, k-core=2, max. Depth=100.

### 2.4 scRNA-seq and Bioinformatic Analysis

- **Tissue acquisition**: Transbronchial lung biopsies (TBLB) from 4 treatment-naïve IPF patients (2023–2024) and 3 controls with benign pulmonary nodules
- **Sample Preparation**:

○ Single-cell suspension (1×10⁵ cells/mL) in PBS (HyClone).
○ Single-cell capture using GEXSCOPE® microfluidic chip.
○ Addition of beads with unique cell barcodes to microwells (1 bead/well).
- **mRNA Capture & Library Construction**:

○ Post-lysis, mRNA is captured by beads tagged with cell barcodes and UMIs.
○ mRNA reverse-transcription to cDNA, followed by amplification.
○ Library construction using GEXSCOPE® Single-Cell RNA Library Kit.
- **Sequencing**:

○ Library diluted to 4 nM.
○ Sequenced on Illumina HiSeq X (150 bp paired-end).
- **Data Analysis**:

○ **Quantification**:

- Group reads/UMIs/genes by cell barcode; count UMIs per gene/cell.
○ **Clustering & Cell Type Identification**:

- Use Seurat (R package, v.3.0.1) for analysis.
- Import expression matrix via read.table; cluster cells using FindClusters (resolution = 0.6).

## 3. Results

### 3.1 Characteristics of Enrolled Subjects

Compared with the control group, the IPF group showed no significant differences in age, gender composition, or smoking history. Among the four parameters of lung function, the percentage of measured to predicted value of (diffusing capacity for carbon monoxide, DLCO) (DLCO%) and (total Lung Capacity, TLC) (TLC%) were significantly lower in the IPF group than in the control group. However, there were no significant differences in the percentage of measured to predicted value of (forced vital capacity, FVC) (FVC%) and (forced expiratory volume in the first second, FEV1) (FEV1%) between the IPF group and the control group. The total number of white blood cells and the absolute value of neutrophils in peripheral blood were significantly higher in the IPF group than in the control group, while the percentage of lymphocytes was significantly lower in the IPF group. Additionally, the levels of cytokeratin 19 fragment and LDH were significantly higher in the IPF group compared to the control group (Table 1).

**Table 1.**
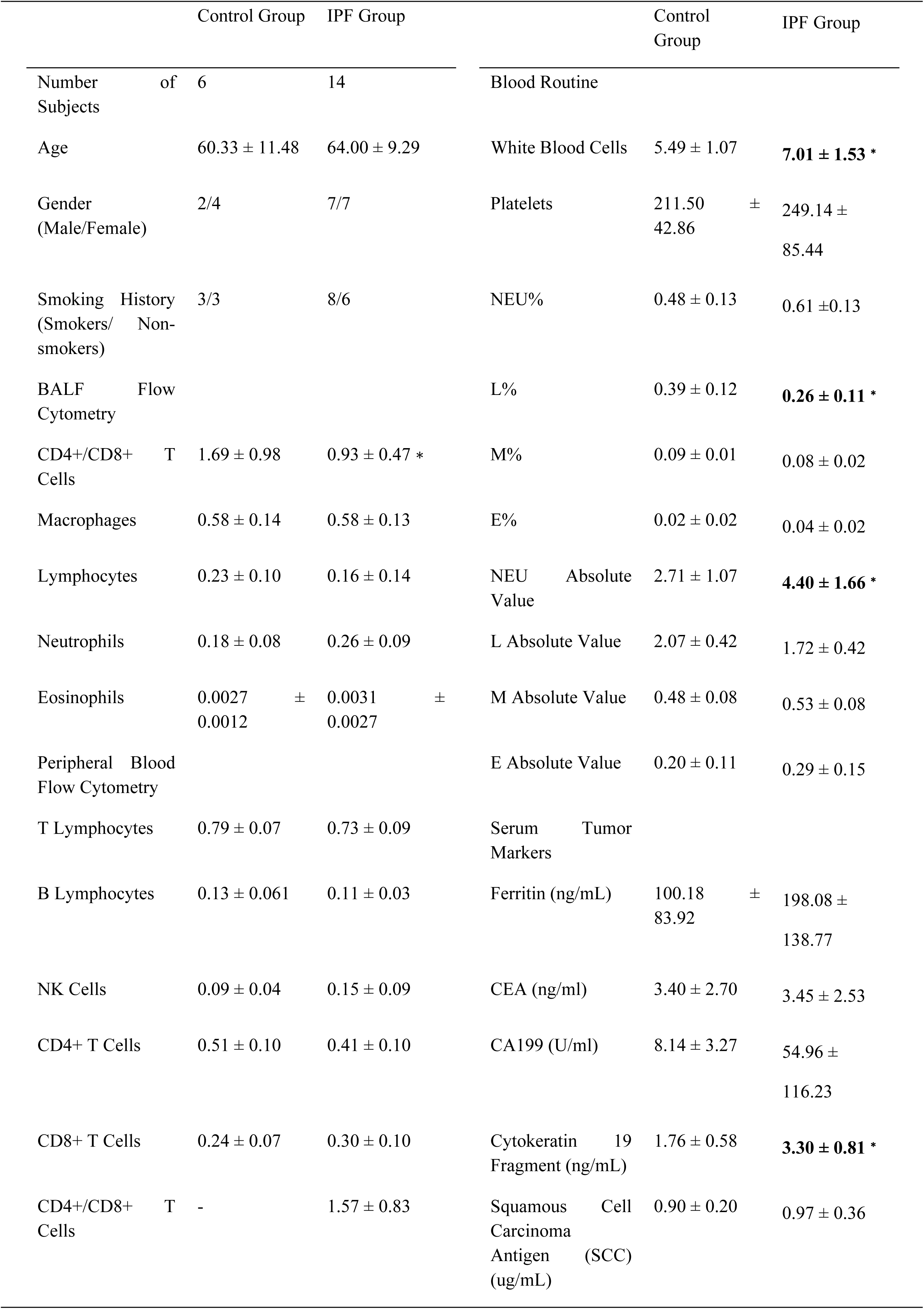
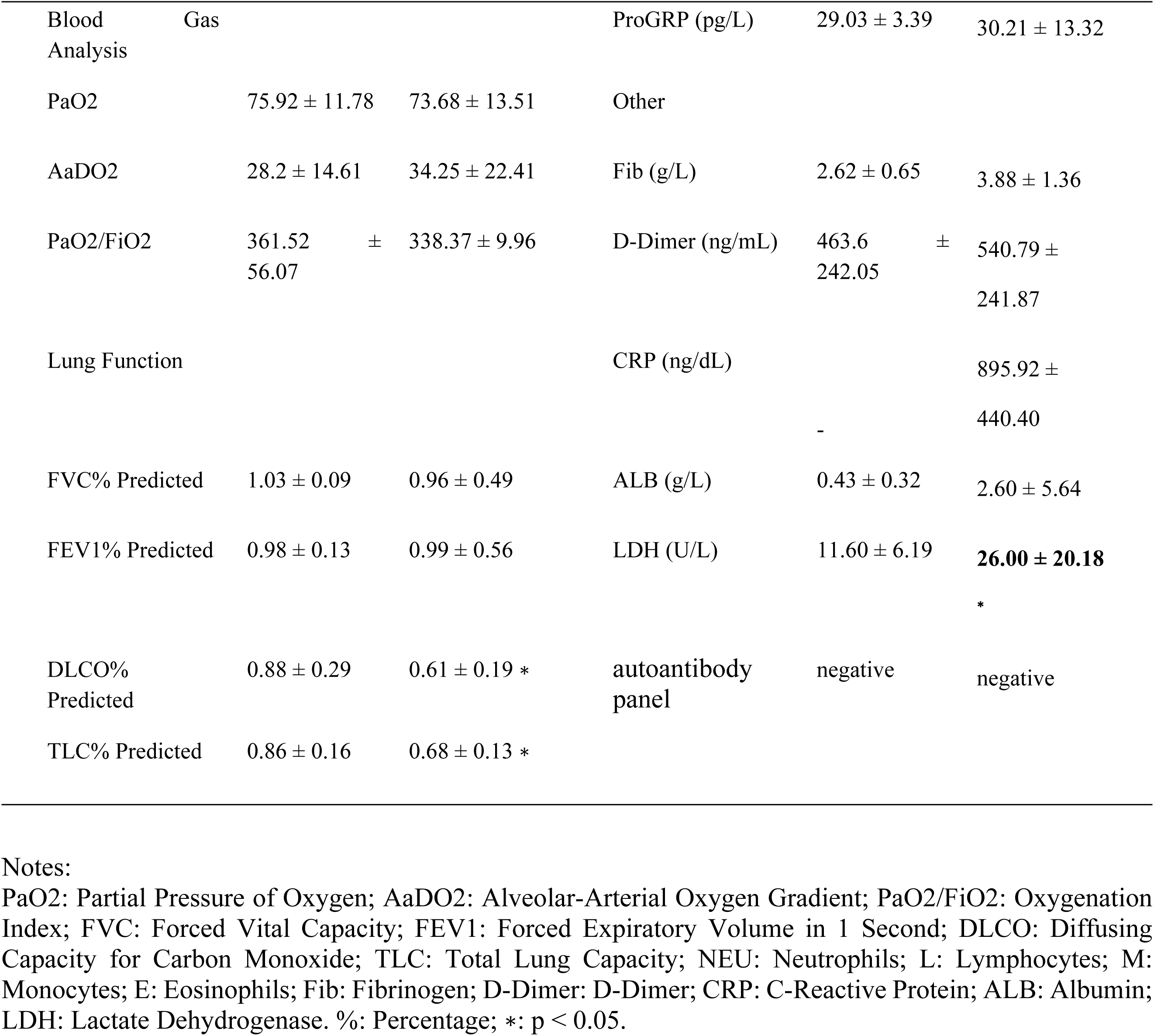
Clinical, Laboratory, and Examination Characteristics of Subjects.

### 3.2 Statistics of DEGs between Groups

Differential expression analysis between the IPF group and the control group revealed 108 DEGs with a fold change >2, including 24 upregulated and 84 downregulated genes (as shown in Table 2). Among the 84 downregulated genes, 4 were unknown genes. The analysis of DEG expression patterns between groups is shown in Figure 2.

**Figure 2.**
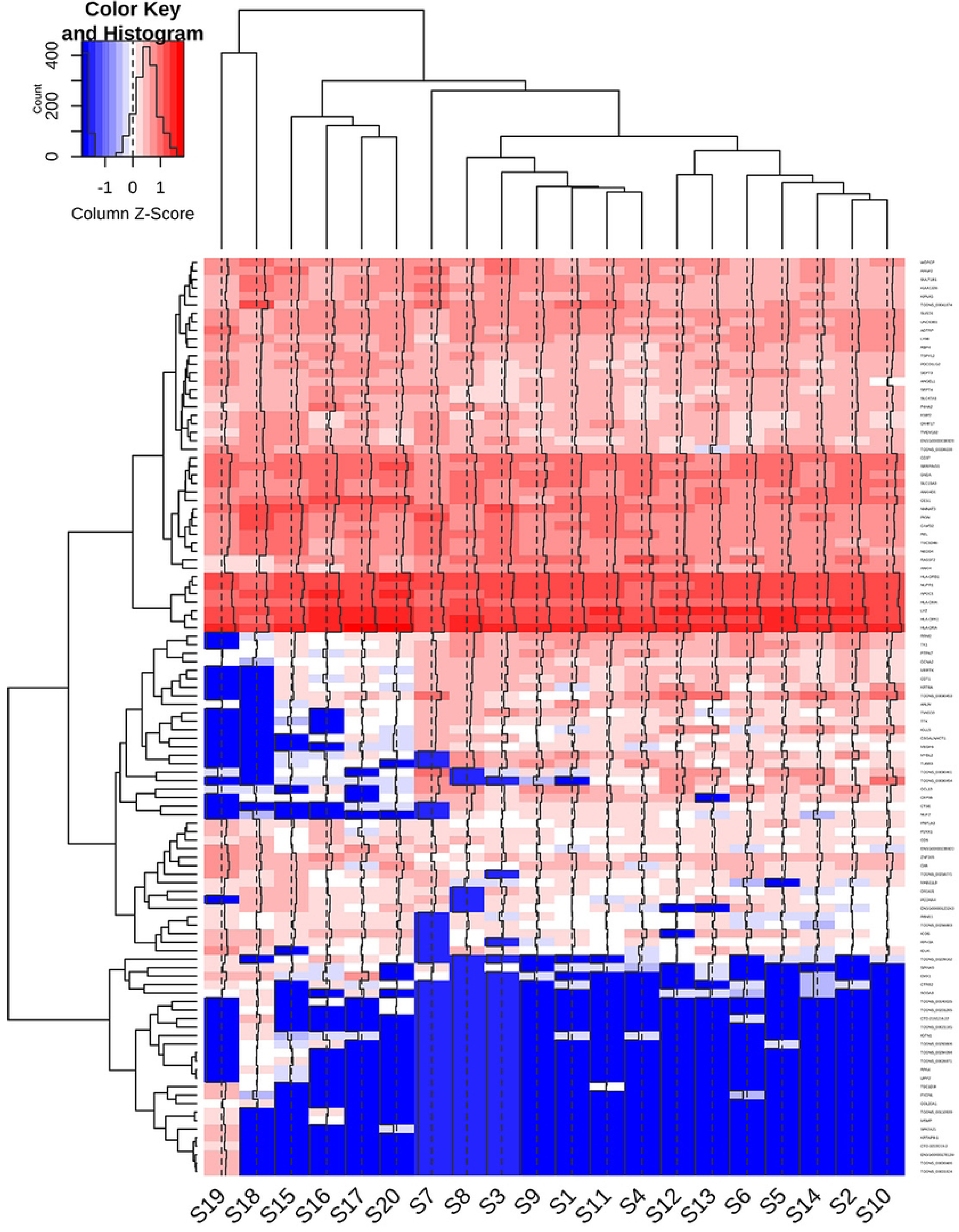
Analysis of DEG Expression Patterns between Groups IPF group: S1-S14; Control group: S15-S20.

**Table 2.** The differentially expressed genes in IPF compared to the control group.

| Gene | Full name/description of gene | IPF fpkm | Control fpkm | FDR | FC |
| --- | --- | --- | --- | --- | --- |
| **A, Upregulated** |  |  |  |  |  |
| CCL13 | C-C motif chemokine ligand 13 | 14.032 | 0.333 | 0.000 | 32.33 |
| NUF2 | NDC80 kinetochore complex component | 0.538 | 0.010 | 0.016 | 29.87 |
| IGLL5 | immunoglobulin lambda like polypeptide 5 | 4.886 | 0.195 | 0.009 | 26.72 |
| TUBB3 | tubulin beta 3 class III | 3.038 | 0.080 | 0.004 | 25.27 |
| MEGF6 | multiple EGF like domains 6 | 0.249 | 0.005 | 0.003 | 22.48 |
| MYBL2 | MYB proto-oncogene like 2 | 1.651 | 0.073 | 0.001 | 20.67 |
| CTSE | cathepsin E | 0.469 | 0.015 | 0.009 | 19.48 |
| TMIGD3 | transmembrane and immunoglobulin domain containing 3 | 5.530 | 0.275 | 0.023 | 14.94 |
| KRT6A | keratin 6A | 1.887 | 0.312 | 0.030 | 12.41 |
| TTK | TTK protein kinase | 0.999 | 0.177 | 0.002 | 11.30 |
| CEP55 | centrosomal protein 55 | 1.130 | 0.093 | 0.016 | 10.11 |
| CSGALNACT1 | chondroitin sulfate N-acetylgalactosaminyltransferase 1 | 0.629 | 0.153 | 0.046 | 8.96 |
| RRM2 | ribonucleotide reductase regulatory subunit M2 | 4.775 | 1.012 | 0.009 | 7.68 |
| ANLN | anillin actin binding protein | 1.000 | 0.358 | 0.031 | 6.58 |
| MERTK | MER proto-oncogene, tyrosine kinase | 1.198 | 0.223 | 0.022 | 6.55 |
| CDT1 | chromatin licensing and DNA replication factor 1 | 1.355 | 0.173 | 0.042 | 6.16 |
| TK1 | thymidine kinase 1 | 4.821 | 0.757 | 0.006 | 6.09 |
| PTPN7 | protein tyrosine phosphatase, non-receptor type 7 | 2.874 | 0.640 | 0.026 | 4.87 |
| CCNA2 | cyclin A2 | 0.771 | 0.135 | 0.009 | 4.75 |
| RASSF2 | Ras association domain family member 2 | 6.725 | 1.783 | 0.005 | 3.61 |
| ANKH | ANKH inorganic pyrophosphate transport regulator | 4.789 | 2.583 | 0.046 | 2.28 |
| TCONS_00090454 | > P01859.2 RecName: Full=Immunoglobulin heavy constant gamma 2 | 7.958 | 0.103 | 0.042 | 53.09 |
| TCONS_00090453 | > P01857.1 RecName: Full=Immunoglobulin heavy constant gamma 1 | 26.494 | 1.390 | 0.042 | 13.21 |
| TCONS_00090461 | > AAS01770.1 monoclonal IgM antibody heavy chain | 9.505 | 0.167 | 0.026 | 31.19 |
| **B, Downregulated** |  |  |  |  |  |
| SUSD1 | sushi domain containing 1 | 4.516 | 7.787 | 0.044 | 0.50 |
| GM2A | GM2 ganglioside activator | 8.196 | 14.097 | 0.036 | 0.49 |
| HLA-DRB1 | major histocompatibility complex, class II, DR beta 1 | 176.759 | 280.055 | 0.036 | 0.49 |
| HLA-DRA | major histocompatibility complex, class II, DR alpha | 534.629 | 907.143 | 0.016 | 0.49 |
| UNC93B1 | unc-93 homolog B1 | 5.762 | 9.622 | 0.030 | 0.48 |
| SLC15A3 | solute carrier family 15 member 3 | 29.314 | 52.350 | 0.045 | 0.48 |
| HLA-DMA | major histocompatibility complex, class II, DM alpha | 75.636 | 132.707 | 0.003 | 0.46 |
| HLA-DPA1 | major histocompatibility complex, class II, DP alpha 1 | 198.577 | 354.827 | 0.003 | 0.45 |
| CD37 | CD37 molecule | 54.187 | 119.858 | 0.045 | 0.45 |
| ANKHD1 | ankyrin repeat and KH domain containing 1 | 6.666 | 12.168 | 0.035 | 0.43 |
| LYZ | lysozyme | 223.384 | 407.753 | 0.001 | 0.42 |
| NUPR1 | nuclear protein 1, transcriptional regulator | 106.514 | 191.955 | 0.003 | 0.42 |
| TSPYL2 | TSPY-like 2 | 2.517 | 5.203 | 0.006 | 0.39 |
| APOC1 | apolipoprotein C1 | 363.112 | 684.093 | 0.031 | 0.39 |
| REL | REL proto-oncogene, NF-kB subunit | 2.621 | 5.562 | 0.049 | 0.36 |
| SLC47A1 | solute carrier family 47 member 1 | 2.221 | 2.900 | 0.035 | 0.35 |
| P4HA2 | prolyl 4-hydroxylase subunit alpha 2 | 2.589 | 5.413 | 0.022 | 0.34 |
| LY86 | lymphocyte antigen 86 | 10.991 | 20.318 | 0.029 | 0.34 |
| RBP4 | retinol binding protein 4 | 9.249 | 21.043 | 0.034 | 0.34 |
| ADTRP | androgen dependent TFPI regulating protein | 8.349 | 20.945 | 0.002 | 0.33 |
| ANGEL1 | angel homolog 1 (Drosophila) | 0.815 | 1.667 | 0.012 | 0.33 |
| Septin-4 | septin 4 | 3.606 | 7.417 | 0.022 | 0.33 |
| Septin-3 | septin 3 | 0.968 | 2.870 | 0.004 | 0.29 |
| SERPING1 | serpin family G member 1 | 21.135 | 57.197 | 0.000 | 0.29 |
| CD6 | CD6 molecule | 0.446 | 1.095 | 0.040 | 0.29 |
| RPAP2 | RNA polymerase II associated protein 2 | 1.671 | 3.752 | 0.031 | 0.29 |
| PDCD1LG2 | programmed cell death 1 ligand 2 | 1.671 | 4.560 | 0.000 | 0.28 |
| ZNF365 | zinc finger protein 365 | 1.936 | 4.788 | 0.033 | 0.28 |
| TMEM182 | transmembrane protein 182 | 6.274 | 23.780 | 0.009 | 0.28 |
| SULT1B1 | sulfotransferase family 1B member 1 | 1.520 | 4.178 | 0.030 | 0.27 |
| KPNA5 | karyopherin subunit alpha 5 | 1.143 | 2.350 | 0.015 | 0.25 |
| WDPCP | WD repeat containing planar cell polarity effector | 4.059 | 10.557 | 0.003 | 0.24 |
| PIGN | phosphatidylinositol glycan anchor biosynthesis class N | 9.861 | 30.345 | 0.009 | 0.21 |
| NEDD4 | neural precursor cell expressed, developmentally down-regulated 4, E3 ubiquitin protein ligase | 24.949 | 78.255 | 0.006 | 0.21 |
| PNPLA3 | patatin like phospholipase domain containing 3 | 0.626 | 2.315 | 0.017 | 0.20 |
| P2RX1 | purinergic receptor P2X 1 | 0.351 | 1.235 | 0.023 | 0.20 |
| ICOS | inducible T-cell costimulator | 0.509 | 2.098 | 0.039 | 0.19 |
| C4orf32 | chromosome 4 open reading frame 32 | 2.940 | 11.572 | 0.004 | 0.19 |
| RBM11 | RNA binding motif protein 11 | 0.494 | 2.267 | 0.011 | 0.19 |
| CES1 | carboxylesterase 1 | 20.467 | 161.992 | 0.000 | 0.18 |
| NMNAT3 | nicotinamide nucleotide adenylyltransferase 3 | 6.244 | 23.363 | 0.002 | 0.17 |
| KIAA1328 | KIAA1328 | 6.859 | 30.417 | 0.002 | 0.17 |
| MAB21L3 | mab-21 like 3 | 0.227 | 0.935 | 0.049 | 0.16 |
| OR14J1 | olfactory receptor family 14 subfamily J member 1 | 0.109 | 0.575 | 0.010 | 0.16 |
| OR4F17 | olfactory receptor family 4 subfamily F member 17 | 8.176 | 37.458 | 0.002 | 0.16 |
| C8B | complement component 8, beta polypeptide | 1.528 | 8.360 | 0.016 | 0.16 |
| PCDHA4 | protocadherin alpha 4 | 0.107 | 0.395 | 0.035 | 0.15 |
| FSIP2 | fibrous sheath interacting protein 2 | 1.668 | 8.237 | 0.000 | 0.15 |
| TBC1D8B | TBC1 domain family member 8B | 5.252 | 21.810 | 0.000 | 0.15 |
| IDUA | iduronidase, alpha-L- | 0.522 | 1.932 | 0.039 | 0.13 |
| RPH3A | rabphilin 3A | 0.166 | 1.400 | 0.001 | 0.09 |
| SPINK9 | serine peptidase inhibitor, Kazal type 9 | 0.281 | 2.695 | 0.046 | 0.07 |
| SOGA3 | SOGA family member 3 | 0.036 | 0.153 | 0.029 | 0.04 |
| IGFN1 | immunoglobulin-like and fibronectin type III domain containing 1 | 0.001 | 0.032 | 0.034 | 0.03 |
| CTRB2 | chymotrypsinogen B2 | 0.040 | 2.482 | 0.000 | 0.02 |
| UPP2 | uridine phosphorylase 2 | 0.001 | 0.345 | 0.042 | 0.02 |
| MSMP | microseminoprotein, prostate associated | 0.001 | 1.153 | 0.046 | 0.02 |
| RPA4 | replication protein A4 | 0.001 | 0.250 | 0.030 | 0.02 |
| TBC1D3I | TBC1 domain family member 3I | 0.008 | 0.717 | 0.035 | 0.01 |
| PXDNL | peroxidasin like | 0.001 | 1.432 | 0.016 | 0.01 |
| COL20A1 | collagen type XX alpha 1 chain | 0.001 | 0.147 | 0.012 | 0.01 |
| KRTAP8-1 | keratin associated protein 8-1 | 0.001 | 1.838 | 0.030 | 0.01 |
| EMX1 | empty spiracles homeobox 1 | 0.052 | 1.562 | 0.000 | 0.01 |
| SPATA21 | spermatogenesis associated 21 | 0.001 | 0.218 | 0.009 | 0.01 |
| CTD-3222D19.2 | - | 0.001 | 0.177 | 0.049 | 0.01 |
| CTD-2192J16.22 | - | 0.008 | 2.277 | 0.016 | 0.01 |
| TCONS_00266883 | > XP_020145385.1 gamma-secretase-activating protein isoform X5 | 0.904 | 3.243 | 0.040 | 0.24 |
| TCONS_00306038 | - | 3.416 | 13.038 | 0.000 | 0.22 |
| TCONS_00041674 | >Q9UHZ2.1 RecName: Full=Metastasis-associated lung adenocarcinoma transcript 1 | 3.134 | 25.753 | 0.000 | 0.08 |
| TCONS_00258771 | > EAW48689.1 hCG1817353 | 0.718 | 2.790 | 0.031 | 0.20 |
| TCONS_00209162 | > EAW78716.1 hCG1813636 | 0.344 | 6.908 | 0.013 | 0.06 |
| TCONS_00026971 | - | 0.001 | 3.165 | 0.042 | 0.02 |
| TCONS_00294094 | > EDL08604.1 mCG1030012 | 0.001 | 1.157 | 0.041 | 0.02 |
| TCONS_00201285 | - | 0.001 | 13.950 | 0.045 | 0.02 |
| TCONS_00283606 | - | 0.038 | 7.328 | 0.006 | 0.01 |
| TCONS_00110939 | > XP_018867249.1 PREDICTED: LOW QUALITY PROTEIN: AFG3-like protein | 0.001 | 3.360 | 0.026 | 0.01 |
| TCONS_00021185 | - | 0.001 | 14.817 | 0.031 | 0.01 |
| TCONS_00140025 | - | 0.001 | 2.758 | 0.003 | 0.01 |

FC, fold change; FDR, adjusted p-value.

### 3.3 Results of Pathway and Functional Enrichment Analysis

GO enrichment analysis of DEGs showed (as in Figure 3) that upregulated DEGs were mainly enriched in biological processes such as cell cycle, organelle organization, mitotic cell cycle process, cell division, protein complex biogenesis, macromolecular complex subunit organization, cellular component assembly, and small molecule metabolic process. Downregulated DEGs were mainly involved in small molecule metabolic process, cellular component assembly, macromolecular complex subunit organization, lipid metabolic process, and cell cycle.

**Figure 3.**
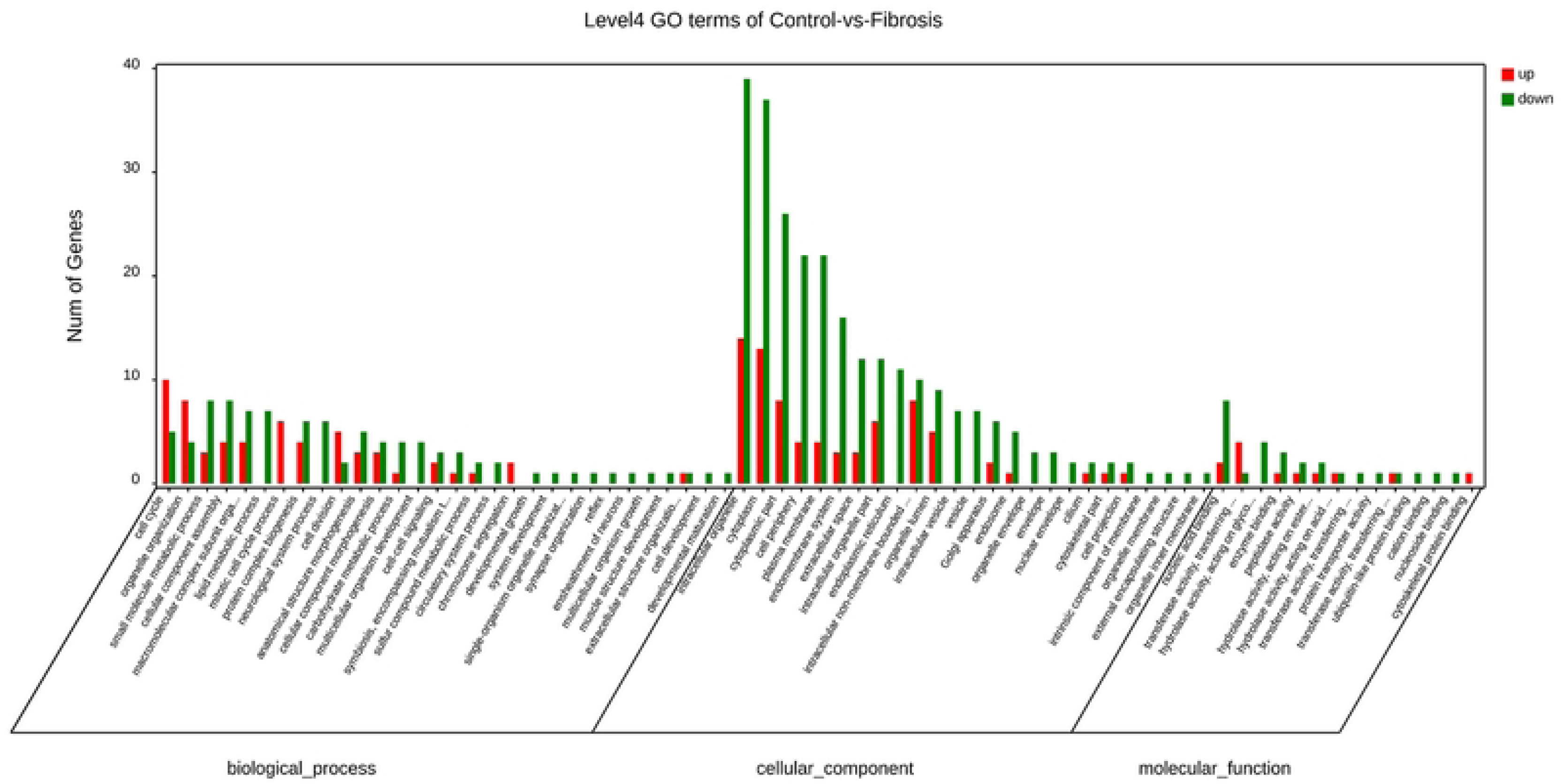
IPF vs Control Group GO Level 4 Classification Bar Chart The x-axis represents GO terms, divided into three major categories from left to right: biological process, cellular component, and molecular function. The y-axis represents the number of genes in the term. Red indicates upregulated genes, and green indicates downregulated genes. GO, Gene Ontology.

The results of KEGG enrichment analysis of DEGs (as shown in Figure 4) revealed that upregulated genes were mainly enriched in inflammation and immune pathways (such as NF-κB signaling pathway, Fc epsilon RI signaling pathway, B cell receptor signaling pathway, phagosome, Fc gamma R-mediated phagocytosis, intestinal immune network for IgA production, natural killer cell-mediated cytotoxicity), pyrimidine metabolism, cell cycle, and PI3K-Akt signaling pathway. Among them, inflammation and immune response seemed to be the most enriched pathways. The most significantly enriched pathway for downregulated DEGs was the cell adhesion molecules (CAMs) pathway, followed by pathways related to immune-related diseases (IgA production intestinal immune network, allograft rejection, systemic lupus erythematosus, autoimmune thyroid disease, and inflammatory bowel disease). These findings highlight the complexity of the immune microenvironment in IPF, which is likely associated with immune dysregulation. This provides new insights for developing therapeutic strategies targeting the immune microenvironment. For example, IgA plays a key role in maintaining immune tolerance in mucosal immunity, including in the respiratory and intestinal tracts. Dysregulation of its production network may indicate impaired mucosal immune tolerance in the lungs of IPF patients, which could be one of the early events in disease pathogenesis. Studies on IgA suggest that it may promote human lung inflammation and fibrosis by enhancing the production of inflammatory or fibrogenic cytokines as well as extracellular matrix, inducing fibroblast differentiation into myofibroblasts, and promoting the proliferation of human lung fibroblasts[12]. This conclusion partially diverges from the findings of our study, and further research is needed to explore the role of these immune-related mechanisms in the pathogenesis of IPF.

**Figure 4.**
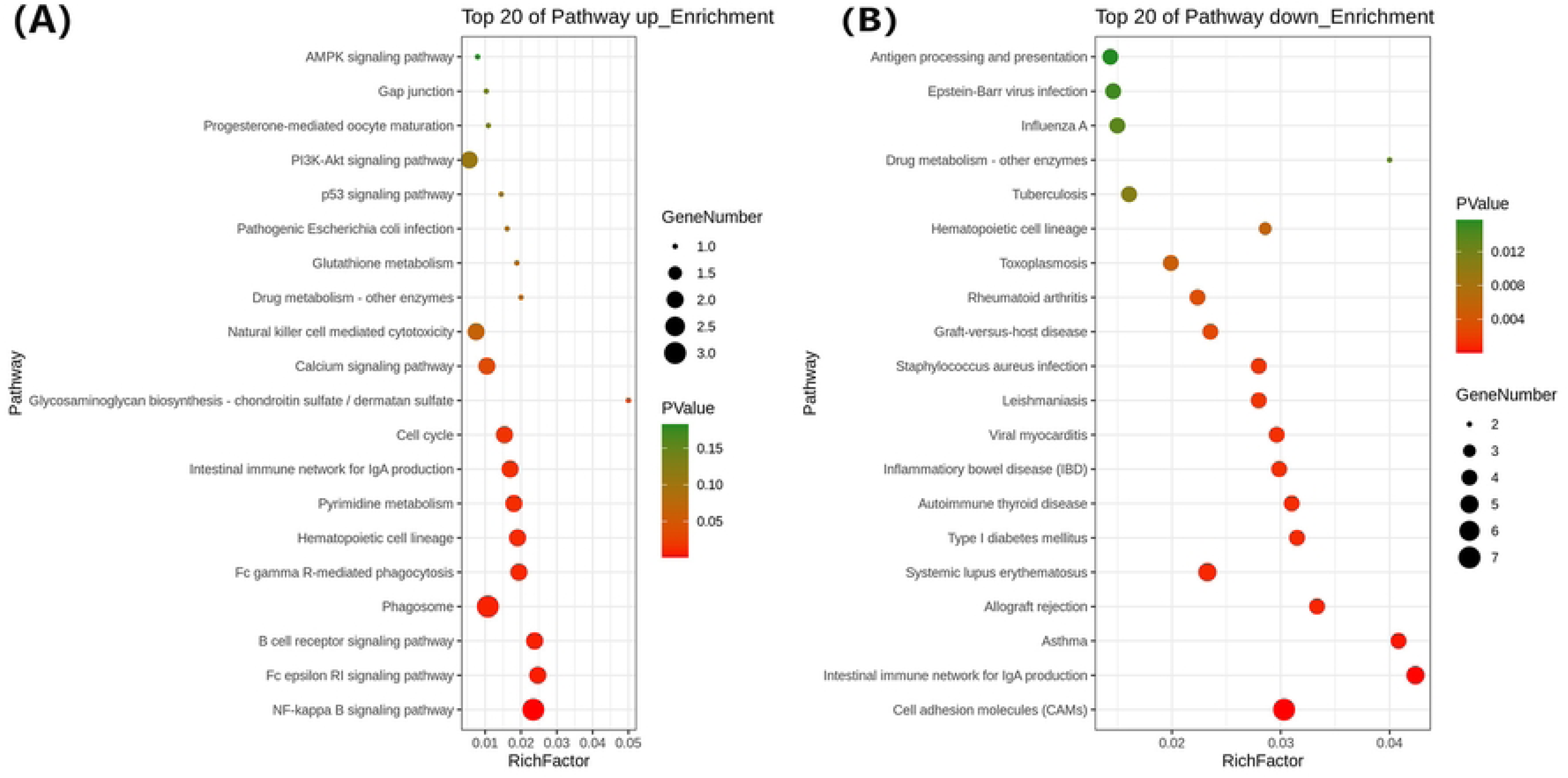
KEGG Enrichment Bubble Chart Showing the top 20 upregulated signaling pathways with the smallest Q value (Figure A) and the top 20 downregulated signaling pathways with the smallest Q value (Figure B). The y-axis represents the signaling pathway name, and the x-axis represents the enrichment factor (RichFactor). GO, Gene Ontology; KEGG, Kyoto Gene and Genome Encyclopedia.

### 3.3 PPI Network Construction and Module Analysis

The 104 DEGs were input into the STRING database to obtain the PPI network diagram. After organizing the PPI network data, it was imported into Cytotype, and the MCODE plugin was used for further clustering and module analysis (parameter settings: degree cutoff=2, node score cutoff=0.2, k-core=2, max. Depth=100). The module with the largest calculated Score value was taken as the final functional module, and three important modules were discovered (as shown in Figure 5), resulting in 17 MCODE genes occupying the most important positions in the PPI network. Among these, nine MCODE genes—NUF2, CEP55, ANLN, TTK, TK1, MYBL2, CCNA2, RRM2, and CDT1—were identified as upregulated (highlighted in red) and constituted Module 1, mainly involved in cell division, cell cycle, and their regulation (Table 3). Eight MCODE genes were downregulated (depicted in blue), with four genes (TBC1D8B, KPNA5, KIAA1328, and RPAP2) forming Module 2, and the remaining four genes (HLA-DRB1, HLA-DMA, HLA-DPA1, and HLA-DRA) constituting Module 3

**Figure 5:**
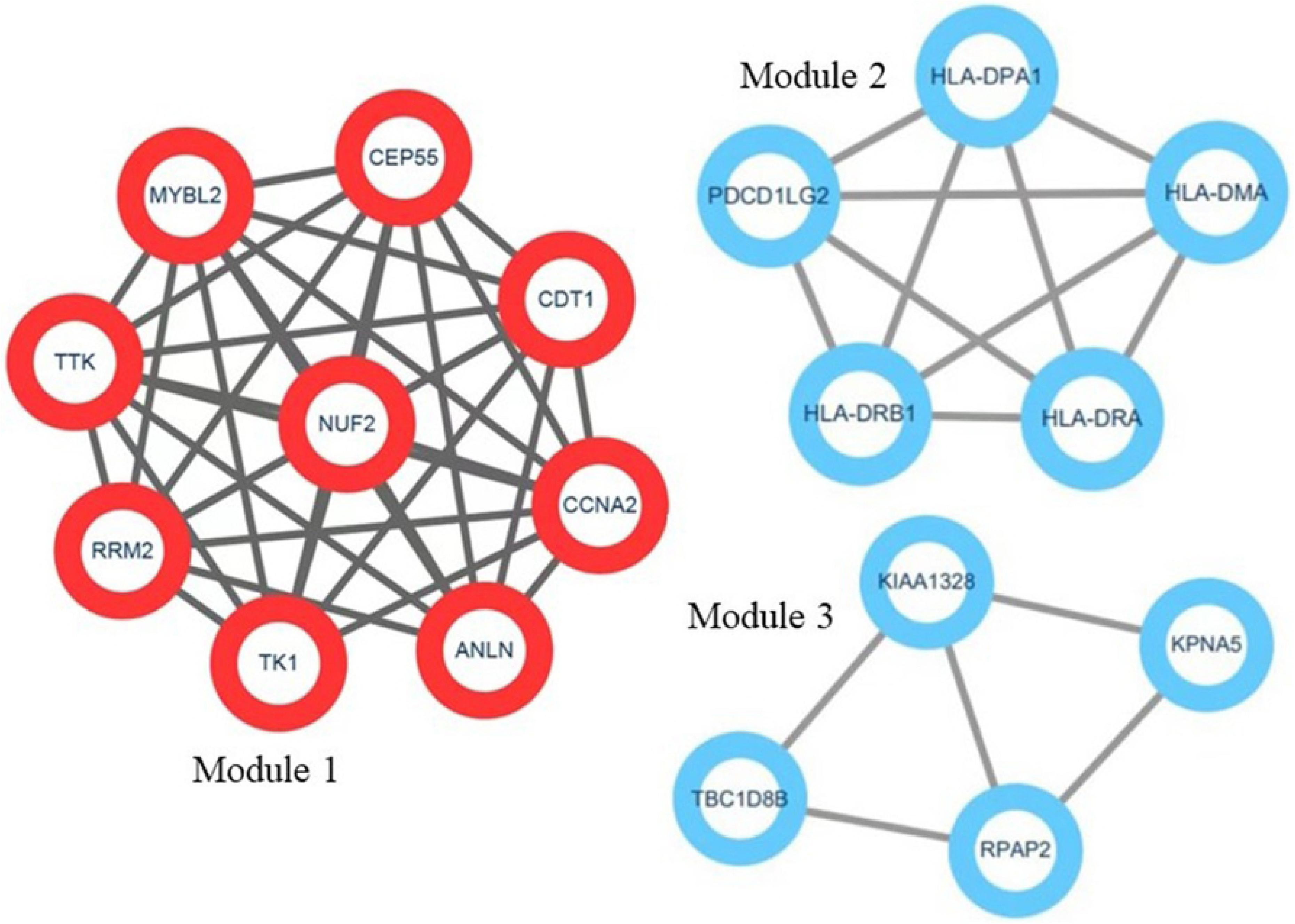
Functional Modules Obtained from MCODE Analysis in Cytoscape. Each node represents a protein, and each line represents the interaction between proteins. Red nodes represent upregulated genes, and blue nodes represent downregulated genes.

**Table 3.** Characteristics of 4 Samples for Single-Cell Sequencing.

| Sample ID | Gender | Age | Race | Smoking | Sampling Method | Sampling Site | Cell Count | Histopathological Result |
| --- | --- | --- | --- | --- | --- | --- | --- | --- |
| IPF1 | Female | 63 | Asia | No | TBLB | Right Upper Lobe | 8099 | UIP |
| IPF2 | Male | 57 | Asia | No | TBLB | Right Lower Lobe | 5601 | UIP |
| IPF3 | Male | 68 | Asia | Yes | TBLB | Right Lower Lobe | 1157 | UIP |
| IPF4 | Male | 67 | Asia | No | TBLB | Right Upper and Middle Lobes | 3683 | UIP |

| Sampl<br>e ID | Gende<br>r | Age | Rac<br>e | Smoking | Sampl<br>ing<br>Metho<br>d | Sampling<br>Site | Cell<br>Cou<br>nt | Histopa<br>thologic<br>al<br>Result |
| --- | --- | --- | --- | --- | --- | --- | --- | --- |
| control<br>1 | Femal<br>e | 55 | Asia | No | TBLB | Right Upper<br>Lobe | 372<br>9 | Normal |
| Contro<br>l2 | Male | 68 | Asia | Yes | TBLB | Right<br>Lower Lobe | 358<br>3 | Normal |
| Contro<br>l3 | Femal<br>e | 30 | Mal<br>e | No | TBLB | Right<br>Lower Lobe | 462<br>5 | Normal |

Note: TBLB refers to Transbronchial Lung Biopsy. UIP refers to Usual Interstitial Pneumonia.

### 3.4 Relationship between IPF Severity and Module 1 Genes

Correlation analysis was performed between the gene expression levels of NUF2, CEP55, ANLN, TTK, TK1, MYBL2, CCNA2, RRM2, and CDT1 in bronchoalveolar lavage fluid (BALF) and the pulmonary function parameters DLCO%, TLC%, FVC%, and FEV1%. This analysis revealed that both NUF2 and TTK expression were negatively correlated with TLC% (Figure 6). Patients were dichotomized into two groups (severe group vs. mild group) based on a threshold of 60% for each of the four pulmonary function parameters. Comparing the gene expression levels of the aforementioned genes in BALF between these groups revealed that NUF2(0.94±.513 VS 0.255±0.205) and ANLN(1.347±0.459 VS 0.51±0.370) expression were significantly higher in the severe group compared to the mild group.

**Figure 6.**
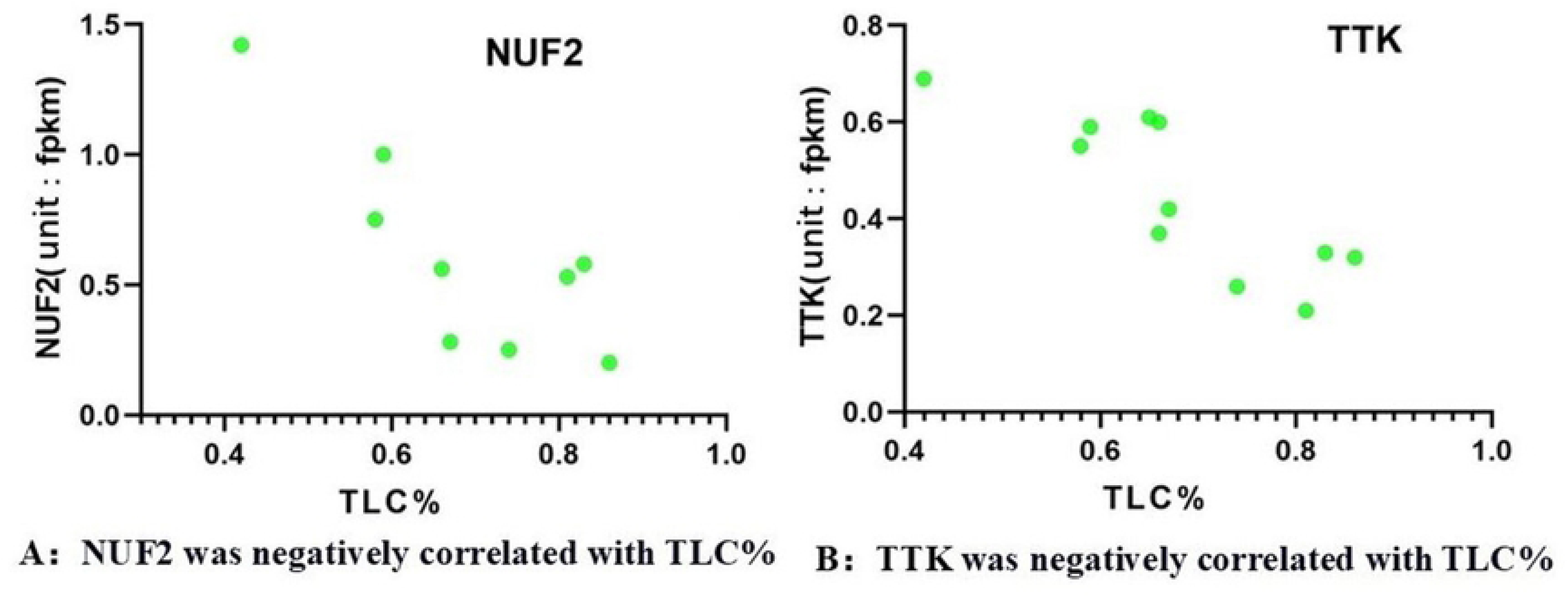
Correlation between TTK and NUF2 with TLC% TTK : TTK protein kinase; NUF2: NDC80 kinetochore complex component; TLC%: the percentage of measured to predicted value of total Lung CapacityTLC%.

### 3.5 Single-Cell RNA Sequencing and Analysis of Target Gene Expression Profiles Across Diverse Cell Types in Lung Tissue

We performed high-resolution single-cell RNA sequencing analysis on 30,477 cells isolated from lung tissues of 4 treatment-naive IPF patients and 3 controls (demographics summarized in Table 4). Unsupervised clustering analysis identified 11 major cell populations (Figure 7): mononuclear macrophages “MPs”, ciliated cells “CiliatedCells”, secretory cells “SecretoryCells”, basal cells “BasalCells”, alveolar epithelial cells “AlveolarEpithelialCells”, endothelial cells “ECs”, proliferating cells “Proliferatingcells”, plasma cells “PlasmaCells”, T cells “TCells”, mast cells “MastCells”, and stromal cells “StromalCells”. Each cell cluster was labeled and identified based on known cell markers (Table 5). Each cell cluster could be further subdivided into subpopulations, and here we mainly highlight the subpopulation data within four pivotal cell types: MPs, ProliferatingCells, StromalCells, and AlveolarEpithelialCells.

**Figure 7.**
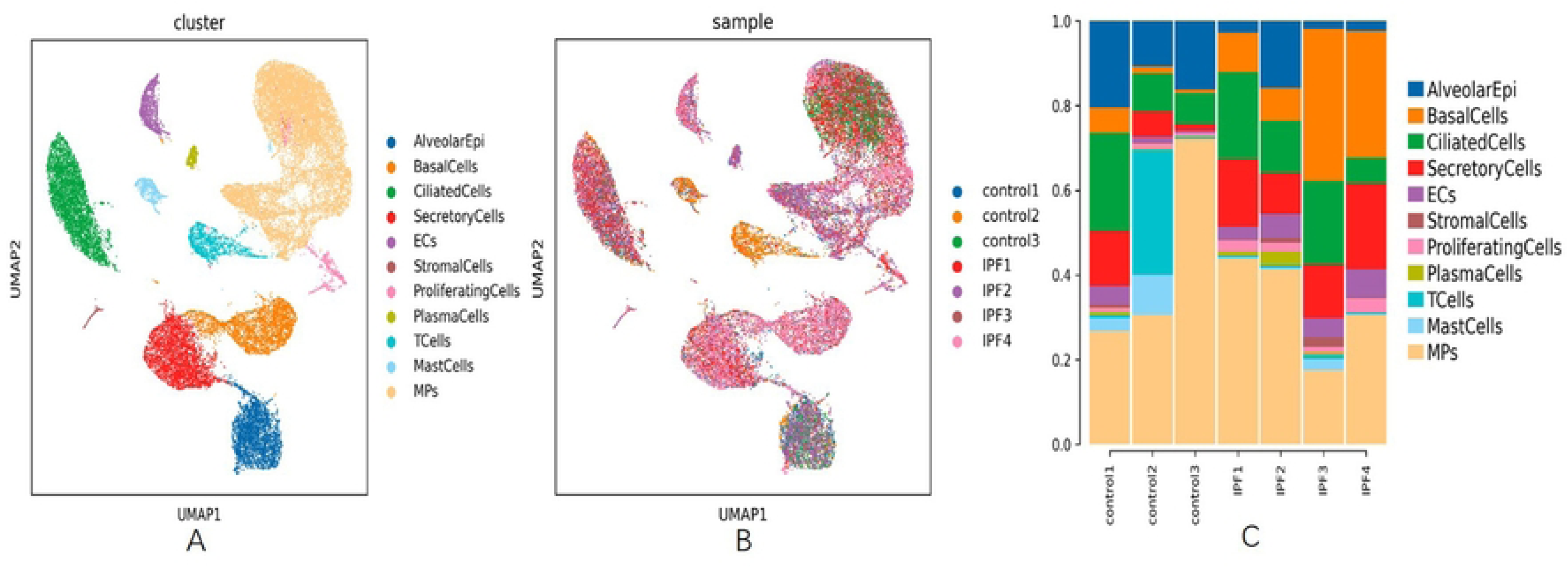
UMAP Plot Colored by Cell Type (A), UMAP Plot Colored by Sample (B), and Bar Chart of Single-Sample Cell Composition (C)

**Table 4:** Cell Type/Abbreviation/Marker Correspondence Table.

| Abbreviation | Cell Type | Marker |
| --- | --- | --- |
| TCells | T Cells | TRBC1, TRBC2, CD2, CD3D |
| MPs | Monocytes/Macrophages | C1QA, MRC1, CD163, CD1C |
| MastCells | Mast Cells | TPSAB1, TPSB2, CPA3 |
| PlasmaCells | Plasma Cells | JCHAIN, MZB1, CD79A, IGKC |
| ECs | Endothelial Cells | CDH5, CLDN5, VWF, PECAM1 |
| CiliatedCells | Ciliated Cells | FOXJ1, PIFO, TPPP3 |
| SecretoryCells | Secretory Cells | WFDC2, MUC5AC, MUC5B, SCGB1A1, SCGB3A1 |
| BasalCells | Basal Cells | KRT15, KRT17, KRT5, KRT19 |
| ProliferatingCells | Proliferating Cells | MKI67, TOP2A, STMN1 |
| AlveolarEpithelialCells | Alveolar Epithelial Cells | SFTPB, SFTPC, ABCA3, LAMP3, CAV1, AGER, CLIC5 |
| StromalCells | Stromal Cells | COL1A1, DCN, LUM, ACTA2, TAGLN |

Note: The table lists various cell types along with their abbreviations and corresponding markers.

As seen in Figure 8, among the upregulated DEGs in IPF compared to the control group, the nine genes (NUF2, CEP55, ANLN, TTK, TK1, MYBL2, CCNA2, RRM2, and CDT1) constituting Module 1 all showed the highest expression levels in “Proliferatingcells”( Figure 8A). The expression profiles of these genes across proliferating cell subclusters are shown in Figure 8B. Next, we computed Module 1 genes signature scores based on Z-scores in each cell type of IPF and control group. Module 1 genes signature demonstrated the highest score in Proliferatingcells Moreover, the Module 1 signature score of Proliferatingcells was significantly higher in IPF than in control group (figure 8C).

**Figure 8.**
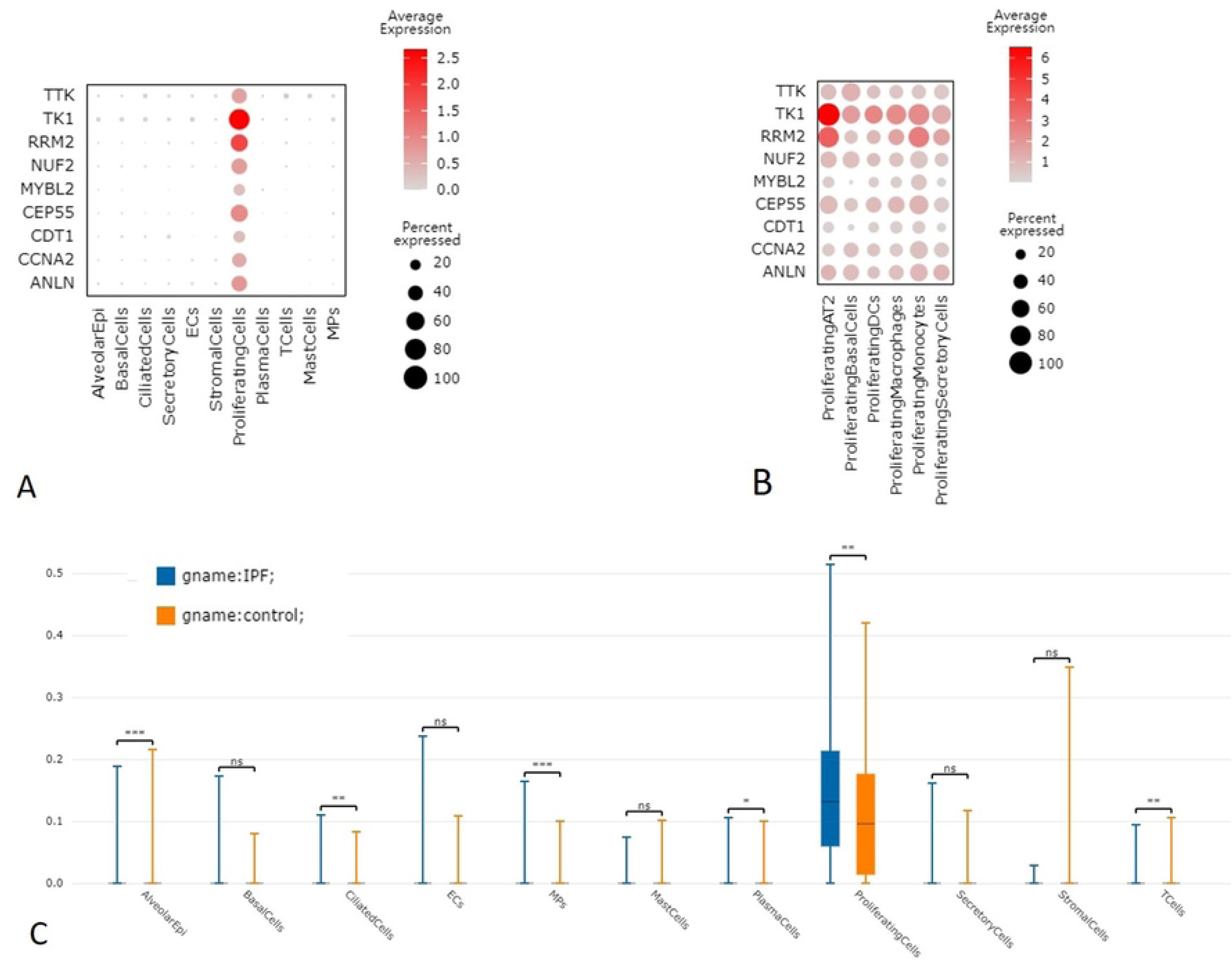
Expression of Module 1 genes in Different Cell Populations(A) and Proliferating Cell Subpopulations(B), and Module 1 genes signature scores(C).

## 4. Discussion

Nine upregulated markers, including NUF2, CEP55, ANLN, TTK, TK1, MYBL2, CCNA2, RRM2, and CDT1, constitute Module 1, which is primarily involved in cell division, cell cycle, and their regulation. Within this module, NUF2 occupies a central position. All nine genes exhibit the highest expression levels in “Proliferating Cells.” Previous research has demonstrated the involvement of these nine genes in the pathogenesis of malignant tumors, with CCNA2 being the only gene studied in the context of IPF ^[13].^

NUF2 is significantly associated with cell proliferation, cell division, and cell cycle arrest. In a rat model of chest irradiation-induced lung cell senescence, NUF2 is significantly upregulated and deemed a central gene. The expression levels of NUF2 in lung adenocarcinoma and lung squamous cell carcinoma are significantly higher than those in normal tissues, and patients with non-small cell lung cancer with high NUF2 expression have a poor prognosis ^[14]^. Studies have also corroborated that NUF2 can augment the expression level of IL8, thereby stimulating the proliferation and survival of CD34+ cells in bone marrow stromal cells, leading to megakaryocyte proliferation and differentiation in primary myelofibrosis ^[15]^. This study found that the expression level of NUF2 in BALF of IPF patients was significantly increased compared with the control group. Single-cell RNA sequencing revealed that NUF2 is predominantly highly expressed in proliferating cells.

TK1 is considered a cell cycle-dependent marker of cell proliferation, reflecting the growth rate of normal cells and various types of tumor cells, with substantial increases observed during DNA synthesis in the S phase of the cell cycle ^[16]^. The TK1 level has been found to be related to tumor proliferation, as TK1 is proportional to the cell proliferation rate ^[17]^. Serum TK1 activity is almost undetectable in healthy individuals but is observed in patients with malignant tumors ^[18]^. Studies have shown that TK1, as a tumor marker, is more sensitive for tumor screening than CEA and AFP ^[19]^. A meta-analysis revealed that TK1 overexpression is associated with a poor prognosis in lung cancer patients ^[15]^. However, no studies have been documented on TK1 in patients with lung fibrosis.

MYBL2, a highly conserved member of the Myb transcription factor family, plays a pivotal role in regulating cell proliferation and survival. MYBL2 is overexpressed in lung adenocarcinoma, and knockdown of MYBL2 can induce apoptosis of lung cancer cells and significantly inhibit cell proliferation, migration, and invasion ^[20]^. Furthermore, upregulated MYBL2 gene expression is a robust predictor of poor prognosis in lung adenocarcinoma ^[21]^.

Chromatin licensing and DNA replication factor 1 (CDT1) are crucial in the cell cycle. In normal cells, precise DNA replication is achieved through licensing of each replication origin once per cell cycle ^[22]^. Licensing is essential for initiating DNA replication at each origin and represents a highly conserved mechanism in eukaryotes ^[23]^. Ectopic expression of CDT1 leads to excessive genome replication and genomic instability in eukaryotes ^[24]^. The expression of CDT1 is downregulated in lung cancer tissues ^[25]^.

CEP55, a mitotic phosphoprotein, plays a vital role in cell division and the cell cycle. During the final stage of cell division, it facilitates the physical separation of two daughter cells ^[26]^. CEP55 is significantly increased in lung cancer cells compared to normal lung tissue ^[27]^. CEP55 overexpression is considered a potential adverse prognostic factor for various cancers, including lung adenocarcinoma ^[28]^, epithelial ovarian cancer ^[29]^, and esophageal squamous cell carcinoma ^[30]^. CEP55 may influence tumor cell proliferation through Myc signaling, DNA repair, and the G2M checkpoint ^[27]^. Beyond its role in cell mitosis and the cell cycle, CEP55 also plays a crucial role in the PI3K/Akt signaling pathway ^[31]^, one of the two main signaling pathways for cell survival/anti-apoptosis. A previous review by our team elaborated on the central role of the imbalance between alveolar epithelial cell apoptosis and fibroblast apoptosis in the pathogenesis of IPF ^[32]^. CEP55 is significantly upregulated in the BALF of IPF patients compared to the control group. Upregulated CEP55 may lead to an imbalance between alveolar epithelial cell apoptosis and fibroblast apoptosis by affecting the cell cycle and the PI3K/AKT signaling pathway, thereby contributing to the pathogenesis and progression of IPF.

ANLN encodes Anillin, an actin-binding protein involved in the PI3K/PTEN signaling pathway ^[33]^. ANLN is upregulated in various cancers ^[34–36]^and has been shown to be a marker of poor prognosis ^[34]^and associated with an aggressive cancer phenotype ^[37]^. Anillin plays an important role in the invasion of breast cancer and pancreatic cancer by regulating the cell cycle ^[38]^. There are no reports on ANLN in IPF or lung fibrosis. This study found that ANLN is significantly upregulated in the BALF of IPF patients.

TTK, a serine, threonine, and tyrosine kinase, has multiple roles in mitosis and meiosis ^[38]^. TTK is deregulated in several human tumors, and its upregulation has been reported in lung cancer, papillary thyroid cancer, breast cancer, gastric cancer, and other cancers, where it is associated with poor prognosis and tumor aggressiveness ^[39]^. Studies have shown that TTK promotes epithelial-to-mesenchymal transition (EMT), a complex cellular process enabling epithelial cells to acquire a mesenchymal phenotype. EMT occurs in three biological contexts: development, wound healing and fibrosis, and tumor progression. Although occurring in three separate biological environments, EMT signaling shares some common molecular mechanisms that allow epithelial cells to dedifferentiate and acquire mesenchymal characteristics, conferring invasive and migratory capabilities to cells toward distant sites. Previous studies have not reported upregulated expression of TTK in IPF patients, but numerous studies have reported that EMT is involved in the fibrotic process of IPF ^[40]^. In our study, TTK was patients, and TTK may participate in the pathogenesis of IPF by promoting EMT.

alveolar epithelium, with most CCNA2-expressing cells also positive for Ki-67 (a proliferation marker) and TUNEL staining in epithelium and hyaline membranes (evidence of apoptosis) ^[13]^. It can be inferred that high CCNA2 expression represents an epithelial proliferative response, and this enhanced proliferation represents a failed compensatory response to injury, indicating that epithelial injury plays an important role in the pathogenesis of AE-IPF. Our study further confirmed that CCNA2 can be detected in the BALF of IPF patients and is significantly upregulated compared to the control group; PPI and module analysis indicated that CCNA2 is involved in Module 1; single-cell RNA sequencing revealed that CCNA2 is predominantly highly expressed in proliferating cells. Consistent with previous studies, it has been shown that CCNA2 mainly originates from proliferating cells.

RRM2 is a rate-limiting enzyme involved in DNA synthesis and damage repair, playing a crucial role in many key cellular processes such as cell proliferation, invasion, migration, and senescence ^[41]^. The expression level of RRM2 is increased in lung cancer tissues compared to normal lung tissues, and high expression levels are associated with poor prognosis ^[42]^. Han et al. found that overexpression of RRM2 promotes the migration and invasion of nasopharyngeal carcinoma cells by inducing EMT. RRM2 can also promote the expression of Bcl-2 ^[43]^. Since Bcl-2 has an anti-apoptotic effect, increased RRM2 expression can upregulate Bcl-2, thereby enhancing the anti-apoptotic effect. Both EMT and fibroblast anti-apoptosis are key mechanisms involved in the pathogenesis of IPF, and RRM2 may participate in the pathogenesis of IPF by affecting EMT and fibroblast apoptosis.

In summary, the expression levels of the nine genes (NUF2, CEP55, ANLN, TTK, TK1, MYBL2, CCNA2, RRM2, and CDT1) are highest in “Proliferating cells.” Previous studies have demonstrated their involvement in the pathogenesis of malignant tumors. Additionally, NUF2 is also associated with lung cell aging, megakaryocyte proliferation, and differentiation^[15]^; CEP55 may also participate in cell anti-apoptosis by affecting the PI3K/AKT signaling pathway ^[31]^; ANLN is also involved in the PI3K/PTEN signaling pathway ^[33]^; TTK and RRM2 promote EMT ^[44,45]^; RRM2 can also upregulate Bcl-2 ^[43]^, leading to increased anti-apoptotic effects; MYBL2 is also involved in regulating cell proliferation and survival ^[20]^. Only CCNA2 has been studied in the context of IPF ^[13]^. In fact, lung cell aging, fibroblast proliferation, alveolar epithelial cell apoptosis, fibroblast anti-apoptosis, EMT, and the PI3K/PTEN signaling pathway are also important mechanisms involved in the pathogenesis of IPF. Although IPF and lung cancer appear unrelated, some scholars have elaborated on the intricate connections between them.

The main limitation of this study is the lack of sufficient experimental validation. Bulk RNA-seq was performed on BALF, which led to the identification of a proliferation-related gene module (Module 1, including NUF2, CEP55, ANLN, TTK, TK1, MYBL2, CCNA2, RRM2, and CDT1). Single-cell RNA-seq was subsequently carried out on lung tissues from a separate cohort of 4 IPF patients and 3 controls. The Module 1 genes, originally derived from BALF, were found to be predominantly expressed in proliferating cells within lung tissues. Moreover, the Module 1 signature score of these proliferating cells was significantly higher in the IPF group than in the control group. This constitutes the sole validation step in our study: markers highly expressed in te BALF of one cohort were corroborated at the transcriptional level in lung tissue samples from another cohort, though both approaches relied solely on sequencing technologies. No additional validation was performed for the proposed markers. Specifically, we did not conduct RT-qPCR to confirm the expression of Module 1 genes in BALF cell pellets or lung tissues. Furthermore, due to the unavailability of remaining lung tissue samples, we were unable to perform experimental validations such as immunohistochemistry, immunofluorescence. To address this limitation, we could obtain specimens via transbronchial lung cryobiopsy in the future. This would allow us to acquire sufficient samples not only for sequencing but also for subsequent validation experiments such as RT-qPCR, immunohistochemistry, and immunofluorescence.

## 5. Conclusion

In the comparison between the IPF group and the control group, bulk RNA-seq was performed on BALF samples, followed by differential expression analysis, which identified 104 DEGs. Among the upregulated markers, NUF2, CEP55, ANLN, TTK, TK1, MYBL2, CCNA2, RRM2, and CDT1 form a functional module primarily involved in cell division, cell cycle regulation, and associated processes. These markers have been more extensively studied in the context of malignant tumors. Single-cell sequencing of cell populations/subpopulations revealed that these nine markers are most highly expressed in proliferating cell populations. This suggests that IPF also exhibits characteristics akin to tumor cell unlimited proliferation. Although the proliferating cells in the lung tissue of IPF patients are not heterogeneous, they may share some pathophysiological mechanisms with tumors, representing a future research avenue. Some targeted antitumor drugs may also be effective for IPF, and further animal experiments are warranted to validate this hypothesis.

## Date availability statement

The datasets generated and/or analyzed during the current study are available in the GEO repository (To review GEO accession GSE312052:Go to https://www.ncbi.nlm.nih.gov/geo/query/acc.cgi?acc=GSE312052 Enter token kjalqsoovrwzzqr into the box).

